# No evidence of coronaviruses or other potentially zoonotic viruses in Sunda pangolins (*Manis javanica*) entering the wildlife trade via Malaysia

**DOI:** 10.1101/2020.06.19.158717

**Authors:** Jimmy Lee, Tom Hughes, Mei-Ho Lee, Hume Field, Jeffrine Japning Rovie-Ryan, Frankie Thomas Sitam, Symphorosa Sipangkui, Senthilvel K.S.S. Nathan, Diana Ramirez, Subbiah Vijay Kumar, Helen Lasimbang, Jonathan H. Epstein, Peter Daszak

## Abstract

The legal and illegal trade in wildlife for food, medicine and other products is a globally significant threat to biodiversity that is also responsible for the emergence of pathogens that threaten human and livestock health and our global economy. Trade in wildlife likely played a role in the origin of COVID-19, and viruses closely related to SARS-CoV-2 have been identified in bats and pangolins, both traded widely. To investigate the possible role of pangolins as a source of potential zoonoses, we collected throat and rectal swabs from 334 Sunda pangolins (*Manis javanica*) confiscated in Peninsular Malaysia and Sabah between August 2009 and March 2019. Total nucleic acid was extracted for viral molecular screening using conventional PCR protocols used to routinely identify known and novel viruses in extensive prior sampling (>50,000 mammals). No sample yielded a positive PCR result for any of the targeted viral families – Coronaviridae, Filoviridae, Flaviviridae, Orthomyxoviridae and Paramyxoviridae. In light of recent reports of coronaviruses including a SARS-CoV-2 related virus in Sunda pangolins in China, the lack of any coronavirus detection in our ‘upstream’ market chain samples suggests that these detections in ‘downstream’ animals more plausibly reflect exposure to infected humans, wildlife or other animals within the wildlife trade network. While confirmatory serologic studies are needed, it is likely that Sunda pangolins are incidental hosts of coronaviruses. Our findings further support the importance of ending the trade in wildlife globally.

## Introduction

The legal and illegal trade in wildlife for consumption as food, medicine and other products is a globally significant threat to conservation (Smith et al., 2006; Nayar, 2009; Rosen and Smith, 2010). It also drives the emergence of pathogens that threaten human and domestic animal health, and national and global economies (Lee and McKibbin, 2004; Smith et al., 2008; Smith et al., 2009). This includes the 2003 Severe Acute Respiratory Syndrome (SARS) outbreak caused by SARS coronavirus (SARS-CoV), which originated in the large wet markets of Guangdong province, China (Ksiazek et al., 2003), and the current COVID-19 outbreak caused by SARS-CoV-2, first discovered in people associated with a wet market in Wuhan (Zhou et al., 2020; Zhu et al., 2020). Both viruses likely originated in bats, with SARS-CoV infecting civets and other small mammals in the markets which may have acted as intermediate or amplifying hosts (Guan et al., 2003; Wang and Eaton, 2007). The finding of furin cleavage insertions in the spike (s) protein sequence in the SARS-CoV-2 genome has led some to suggest that intermediate hosts may have been involved in the emergence of COVID-19 (Andersen et al., 2020), however no intermediate hosts have so far been conclusively identified. Recently, four different groups have identified coronaviruses in imported Sunda or Malayan pangolins (*Manis javanica*) seized in raids on wildlife traders in China (Liu et al., 2019; Lam et al., 2020; Xiao et al., 2020; Zhang et al., 2020). The genomes of these are closely related to SARS-CoV-2, particularly in some genes, including the s-gene responsible for binding to host cells, albeit that some bat-CoVs have higher overall sequence identity to SARS-CoV-2 (Latinne et al., 2020). Authors of these papers propose that further sampling of pangolins might help elucidate the potential role of pangolins in the evolution of SARSr-CoVs, the emergence of COVID-19, and the risk of future zoonotic viral emergence (Liu et al., 2020; Lam et al., 2020; Xiao et al., 2020).

Over a ten-year period, as part of the USAID PREDICT project (PREDICT Consortium, 2017; PREDICT Consortium, 2019), we collected biological samples from confiscated and rescued Sunda pangolins in their country-of-origin: Peninsular Malaysia and the Malaysian state of Sabah on the island of Borneo. The aims of this study were to identify the phylogeographic origins of confiscated pangolins and any potentially zoonotic pathogens associated with them (Karesh 2010). Here we report on the results from pathogen surveillance and discovery screening of these pangolin samples.

## Materials & methods

Sunda pangolins (*Manis javanica*) were either confiscated from smugglers or rescued from the wild between August 2009 and March 2019, and were in the possession of the Department of Wildlife and National Parks Peninsular Malaysia, or Sabah Wildlife Department at the time of sampling. Most confiscations occurred near national borders or ports, and were reported to be destined for other Southeast Asian countries en route to China, and were usually found in sacks or crates in temporary holding facilities, or in vehicles. The wild-rescued Sunda pangolins were all surrendered by members of the public who found them in their native habitats. All pangolins were alive during the sampling process, based on their weakened condition the animals had been in captivity for varying lengths of time, but usually many weeks or months, there was no way of confirming this.

The sampling protocol was approved by UC DAVIS Institutional Animal Care and Use Committee (protocol number: 16048). Each pangolin was assigned a unique identification code, GPS coordinates of the confiscation or rescue locations, biometric measurement and physical health check information were recorded. Swab samples were collected from the throat and rectum using a sterile non-absorbent mini-tip polyester swab (Puritan, Guilford, USA) placed in a cryotube contained 600 μL of TRIzol reagent (Invitrogen, Carlsbad, USA). All samples were stored immediately in a liquid nitrogen dewar MVE Doble 34 (Chart Biomed, Ball Ground, USA) at the sampling site and transferred to a −80°C freezer for long term storage.

Total nucleic acid was extracted for viral molecular screening using the NUCLISENS EASYMAG or MINIMAG system according to the manufacturer’s protocol with validated modifications (bioMérieux, Marcy l’Etoile, France). Complementary DNA (cDNA) of each sample were generated, according to manufacturer’s protocol with random hexamers, from the SuperScript III First-Strand Synthesis System for reverse transcription PCR (Invitrogen, Carlsbad, USA). The cDNA was used in conventional PCR protocols screening five viral families: Coronaviridae, Filoviridae, Flaviviridae, Orthomyxoviridae and Paramyxoviridae (Table 1). The PCRs were conducted in a Veriti or SimpliAmp thermal cycler (Applied Biosystems, Foster City, USA). Reactions were carried out in a final volume of 20 µl, following the manufacturer’s protocol (Qiagen, Hilden, Germany) using 1 µl of the cDNA product as template and either Fast Cycling PCR kit or HotStarTaq *Plus* Master Mix with a final concentration of 0.1 µM for each primer following the manufacturer’s protocols (Qiagen, Hilden, Germany). PREDICT universal controls 1 and 2 (Anthony et al., 2013), and specific controls for Filovirus (One Health Institute Laboratory, University of California, Davis) and Influenza Liang PCR (Liang et al., unpublished) were used. Peninsular Malaysia and Sabah samples were screened on separate occasions at two different certified BSL2 biocontainment level laboratories using standardised methods. PCR products were loaded and run on 1% agarose gel electrophoresis - 100V, for 30-45 minutes with 0.5x tris-acetate-EDTA buffer (Vivantis Technologies Sdn. Bhd., Subang Jaya, Malaysia). The gels were viewed on a transilluminator and expected size bands were excised, stored in separate microcentrifuge tubes, and the corresponding post PCR mixes were used as a template for contamination control PCRs to check for contamination from the universal positive controls. PCR products were run under the same gel electrophoresis conditions; those without the expected size bands showed that there was no contamination from the controls. Products from the initial PCR of these samples were then purified using the Ultrafree-DA centrifugal filter units (Millipore, Cork, Ireland); the purified products were cloned using the dual-colour selection Strataclone PCR cloning kit according to the manufacturer’s protocol (Stratagene, La Jolla, USA). Up to eight colonies containing the PCR product were selected and inoculated on Luria Bertani agar slants, individually. Grown colonies were sent to a commercial company for direct colony sequencing.

**Table 1.**
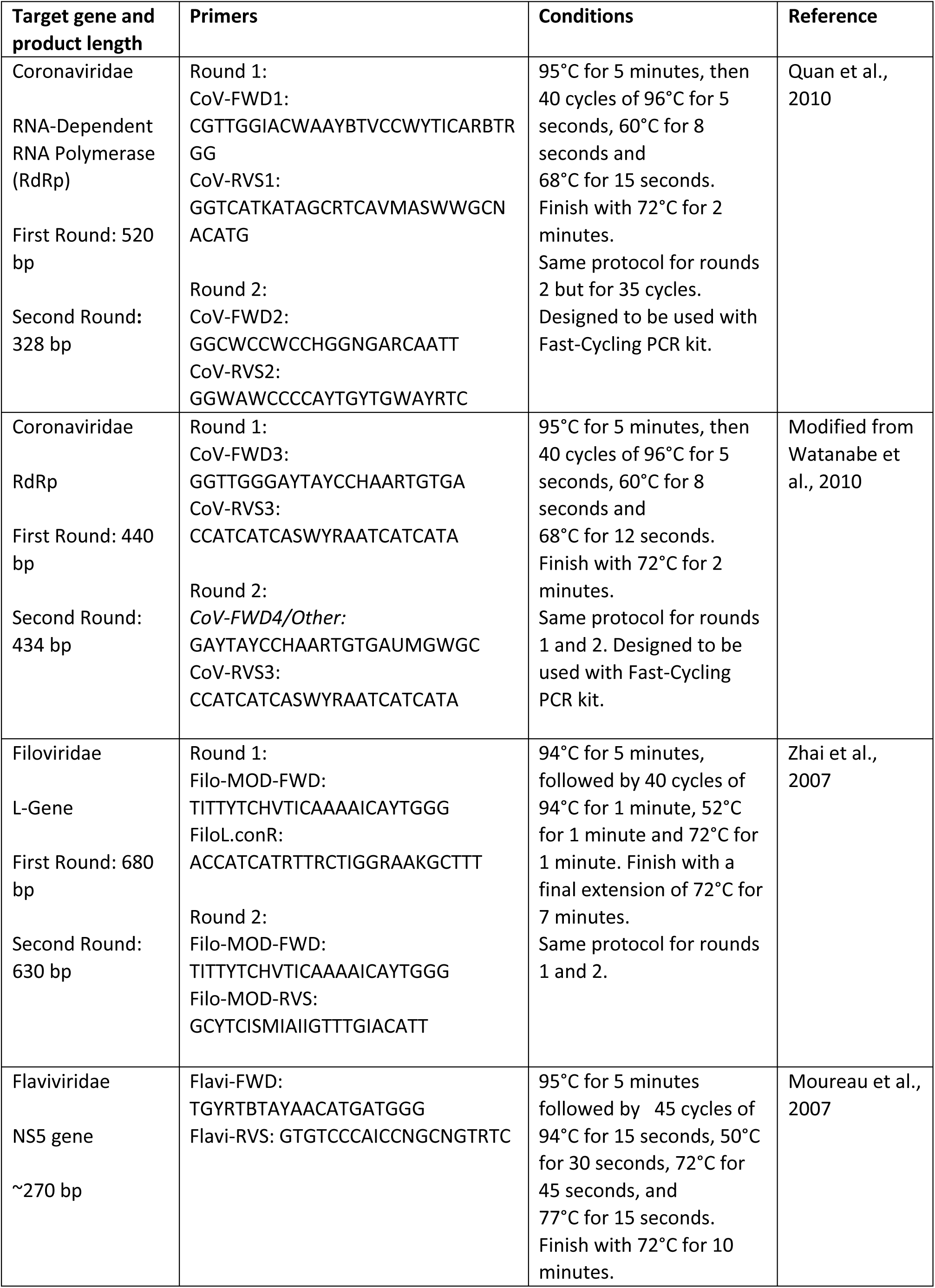
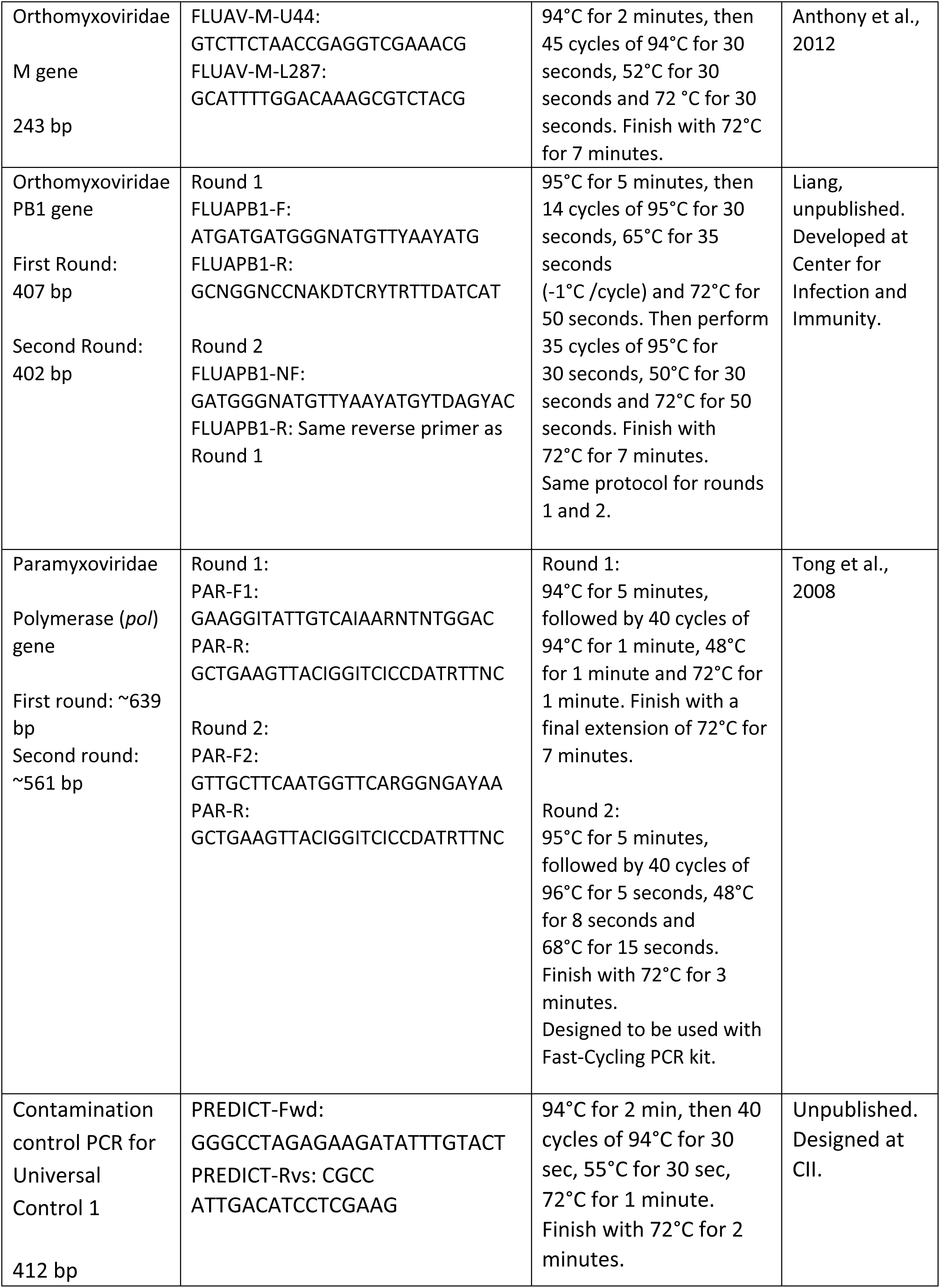
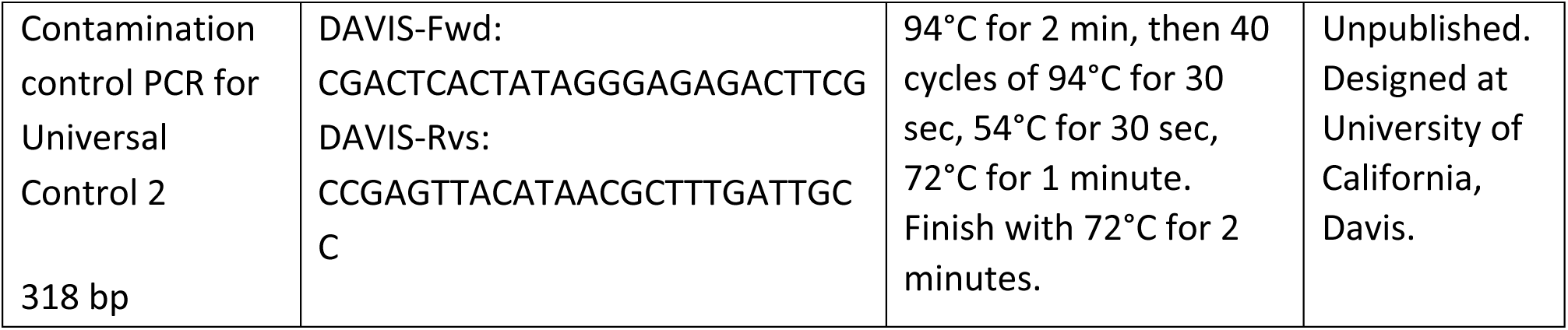
PCR conditions and primer sequences used.

## Results

A total of 334 Sunda pangolins were screened: 289 in Peninsular Malaysia (confiscated n=286; wild-rescued n=3) (Tables 2a and 2b), and 45 in Sabah state (confiscated n=40; wild-rescued n=5) (Tables 2c and 2d). No sample yielded a positive PCR result for any member of the targeted virus families, either in Peninsular Malaysia (95% CI 0.0-0.01) or in Sabah (95% CI 0.0-0.08). All positive controls were successfully amplified, confirming that the PCRs were performing properly.

**Table 2a.**
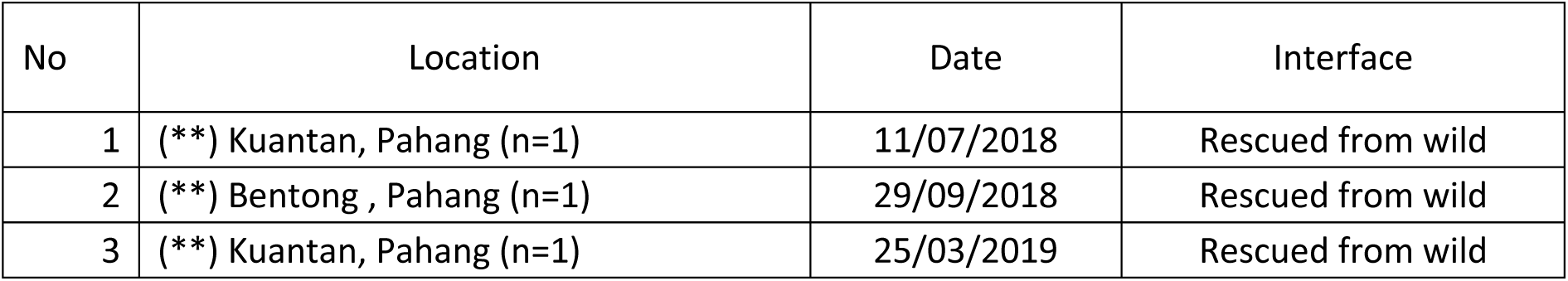
Details of Sunda pangolins rescued from the wild in Peninsular Malaysia

**Table 2b.** Details of Sunda pangolins confiscated from Peninsular Malaysia

| No | Location | Date | Interface |
| --- | --- | --- | --- |
| 1 | (*) Kedah (n=65) | 27/08/2009 | Confiscation |
| 2 | (*) Kelantan (n=73) | 09/09/2009 | Confiscation |
| 3 | (*) Johor (n=18) | 30/09/2009 | Confiscation |
| 4 | (*) Johor (n=2) | 06/11/2009 | Confiscation |
| 5 | (*) Johor (n=40) | 07/11/2009 | Confiscation |
| 6 | (*) Johor (n=45) | 06/04/2010 | Confiscation |
| 7 | (*) Pulau Pinang (n=30) | 27/04/2010 | Confiscation |
| 8 | (*) Johor (n=13) | 22/07/2010 | Confiscation |

**Table 2c.** Details of Sunda pangolins rescued from the wild in Sabah state

| No | Location | Date | Interfaces |
| --- | --- | --- | --- |
| 1 | (**) Tambunan (n=1) | 17/06/2015 | Rescued from wild |
| 2 | (**) Kota Kinabalu (n=1) | 01/05/2016 | Rescued from wild |
| 3 | (**) Penampang (n=1) | 07/06/2016 | Rescued from wild |
| 4 | (**) Kota Kinabalu (n=1) | 21/02/2017 | Rescued from wild |
| 5 | (**) Penampang (n=1) | 18/01/2018 | Rescued from wild |

**Table 2d.** Details of Sunda pangolins confiscated from Sabah state

| No | Location | Date | Interfaces |
| --- | --- | --- | --- |
| 1 | (**) Beaufort (n=10) | 31/10/2014 | Confiscation |
| 2 | (**) Kota Belud (n=2) | 18/09/2015 | Confiscation |
| 3 | (**) Lahad Datu (n=23) | 22/02/2016 | Confiscation |
| 4 | (**) Penampang (n=2) | 29/11/2016 | Confiscation |
| 5 | (**) Sandakan (n=3) | 26/09/2017 | Confiscation |

Tables (2a), (2b), (2c) and (2d) Details of Sunda pangolins confiscated from smugglers and rescued from wild. (Level of detail for the location of confiscations and rescue events reported was determined by the local wildlife departments).
(n) indicates the total number of pangolins sampled in the event,
(\*) indicates the State where the confiscation or rescue occurred in Peninsular Malaysia,
(\*\*) indicates the District where the confiscation or rescue occurred in Peninsular Malaysia or Sabah state.

## Discussion

Our negative findings across five viral families associated with emerging and re-emerging zoonotic diseases in recent decades contrast with reports of the detection of parainfluenza virus (Wang et al., 2018), coronaviruses and Sendai virus (Liu et al., 2019; Zhang et al., 2020), and SARSr-CoVs (Lam et al., 2020; Xiao et al., 2020) in Sunda pangolins. Our sample size is substantial, particularly given the rarity of these animals in Malaysia - the International Union for Conservation of Nature (IUCN) lists the Sunda pangolin (*Manis javanica*) as ‘Critically Endangered’, as a result of poaching, smuggling and habitat loss (IUCN, 2018). Our previous studies of bat coronaviruses revealed 5-10% PCR prevalence (Yang et al., 2016; Anthony et al., 2017; Hu et al., 2017; Latinne et al., 2020), suggesting that even at the upper limit of the 95% confidence interval, our negative findings in pangolins are inconsistent with endemic coronavirus infection at a population level. Serologic studies are needed to support this contention.

While our sampling was necessarily opportunistic (given the conservation status and the cryptic nature of the species) and sampling intensity varied, our negative findings over ten years and at multiple locations supports the veracity of the findings. The most parsimonious explanation for the contrast between our findings and the discovery of SARSr-CoVs in Sunda pangolins by (Liu et al., 2019; Liu et al., 2020; Lam et al., 2020, Xiao et al., 2020, Zhang et al., 2020) is the nature of the sampled population: our samples were drawn from an ‘upstream’ cohort of animals yet to enter or just entering the illegal trade network, whereas all others were drawn from ‘downstream’ cohorts confiscated at their destination in China. During the wildlife trade transits, which often includes movement through other Southeast Asian countries, animals are often housed together in groups from disparate geographic regions, and often with other species, giving opportunity for viral transmission among and within species. The housing of some of the animals in rehabilitation centers in China would also allow for exposure to coronaviruses from other groups or species. In natural wildlife reservoir hosts, SARSr-CoVs appear to cause little if any clinical signs, and this is supported by the limited laboratory infections so far carried out (Watanabe et al., 2010). The reports of clinical illness and pathology associated with coronavirus infection in pangolins (Liu et al., 2019; Xiao et al., 2020), are unlikely in a reservoir host. We therefore conclude that the detections of SARS-CoV-2 related viruses in pangolins are more plausibly a result of their exposure to infected people, wildlife or other animals after they entered the trade network. Thus, the likelihood is that Sunda pangolins are incidental rather than reservoir hosts of coronaviruses as claimed by Zhang et al., (2020).

Our microsatellite DNA fragment analysis (manuscript in preparation) suggests that confiscated pangolins from Peninsular Malaysia and Sabah were taken from Malaysia, Brunei or Indonesia, however further analysis of pangolins from the neighbouring countries is required to confirm the results. They were confiscated at holding facilities, ports or borders prior to shipment, and had not yet been exposed to multiple potential sources of infection, unlike the confiscated animals in China reported by Xiao et al., (2020) and Lam et al., (2020). An array of pathogens and infections have been observed in wet markets, in wildlife (Dong et al., 2007; Cantlay et al., 2017), in humans (Xu et al., 2004) and in domestic animals (Karesh et al., 2005). In comparison to wildlife screened from the wild (Poon et al., 2004) and from farms (Tu et al., 2004; Kan et al., 2005), wildlife in markets have a much higher chance of exposure to pathogens and disease spillover. These findings highlight the importance of carefully and systemically ending the trade in wildlife and improving biosecurity to avoid having wet markets where wild animals are mixing with farmed animals and humans.

Our findings suggest that pangolins that have not entered the illegal wildlife trade pose no threat to human health. While the detection of SARS-CoV-2 like viruses in some trade-rescued pangolins suggests a parallel with traded civets (*Parguma larvata*) in the emergence of SARS-CoV (Guan et al., 2003), any role as an intermediate host in the transmission of SARS-CoV-2 from a putative natural bat host to humans is yet to be established. Serological studies in pre-trade pangolins will shed further light on any role of pangolins as hosts of SARS CoV2-related viruses. All pangolin species face known and significant threats to their survival in nature and require active conservation efforts to ensure their enduring existence for future generations.

## Acknowledgments

We thank the Malaysian Government, particularly the Wildlife Disease Surveillance Programme of the Department of Wildlife and National Parks Peninsular Malaysia, Ministry of Health Malaysia, Department of Veterinary Services Malaysia, Sabah State Health Department, Universiti Malaysia Sabah, Sabah Wildlife Department, Sabah Wildlife Health Unit and Sabah Wildlife Rescue Unit. We thank Dato’ Abdul K.A. Hashim, Rahmat Topani, Augustine Tuuga and Jum R.A. Sukor for their permission and support to conduct this research. We thank members of the EcoHealth Alliance Malaysia field team (Dr. Zahidah Zeid, Mohamed S.M. Azian, Alexter Japrin, Ronald H.M. Tinggu, Muhammad Y. Wazlan, Nor A. Aziz) and Sabah Wildlife Rescue Unit and Wildlife Health Unit (Andrew Ginsos, Runie David and Leonorius bin Lojivis) for assistance with sample collection, the laboratory team (Suraya Hamid, Nur A.M. Sungif) for sample processing at the Molecular Diagnostic Laboratory, at the Department of Wildlife and National Parks Peninsular Malaysia’s National Wildlife Forensic Laboratory and the laboratory team (Fernandes Opook, Emilly Sion) for sample processing at Sabah Wildlife Department’s Wildlife Health, Genetic and Forensic Laboratory and Program Assistant Velsri Sharminie for generating the maps. We dedicate this paper to Dr. Diana Ramirez, who sadly passed away on 31/10/2018, in honour of her vital contribution to this work and wildlife conservation in Sabah. This study was made possible in part by the generous support of the American people through the United States Agency for International Development (USAID) Emerging Pandemic Threats PREDICT project (Cooperative Agreement Numbers AID-OAA-A-14-00102 and GHN-AOO-09-00010-00), and the USAID Infectious Disease Emergence and Economics of Altered Landscapes (IDEEAL) Project (Cooperative Agreement number AID-486-A-13-00005). The contents are the responsibility of the authors and do not necessarily reflect the views of USAID or the United States Government.

---

**Figure 1.1:**
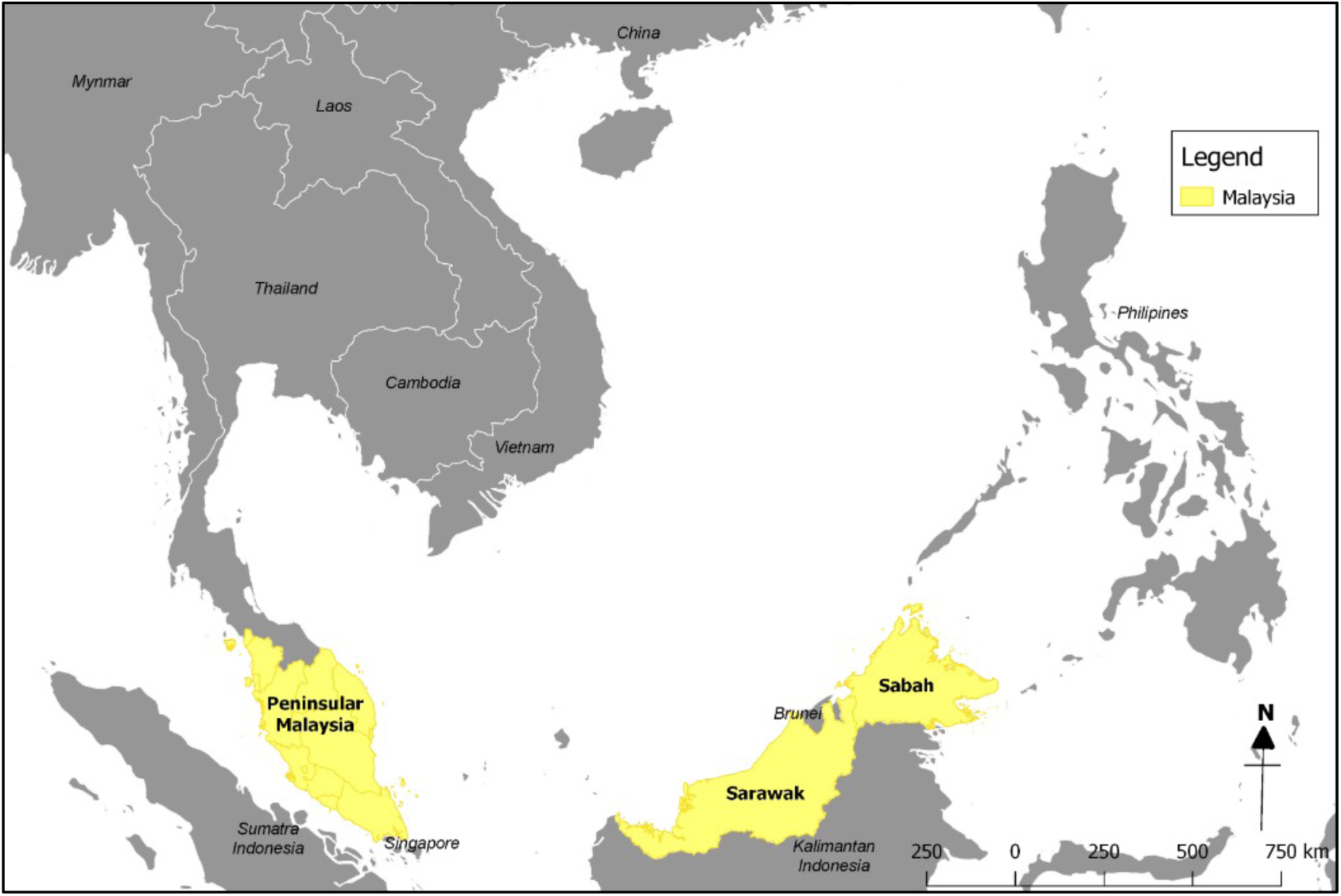
Map of Southeast Asia and China.

**Figure 1.2:**
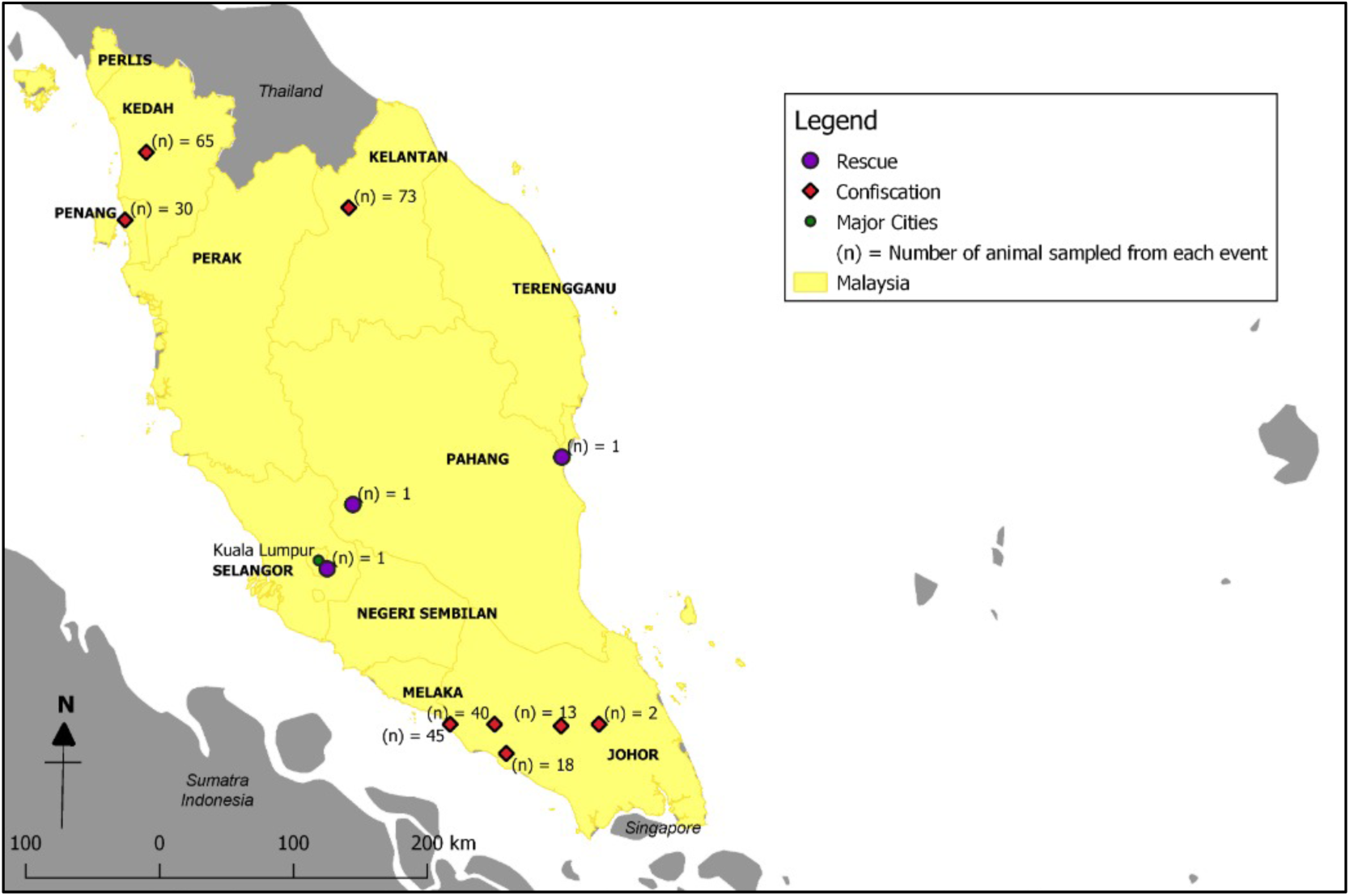
Locations where pangolins were rescued and confiscated in Peninsular Malaysia.

**Figure 1.3:**
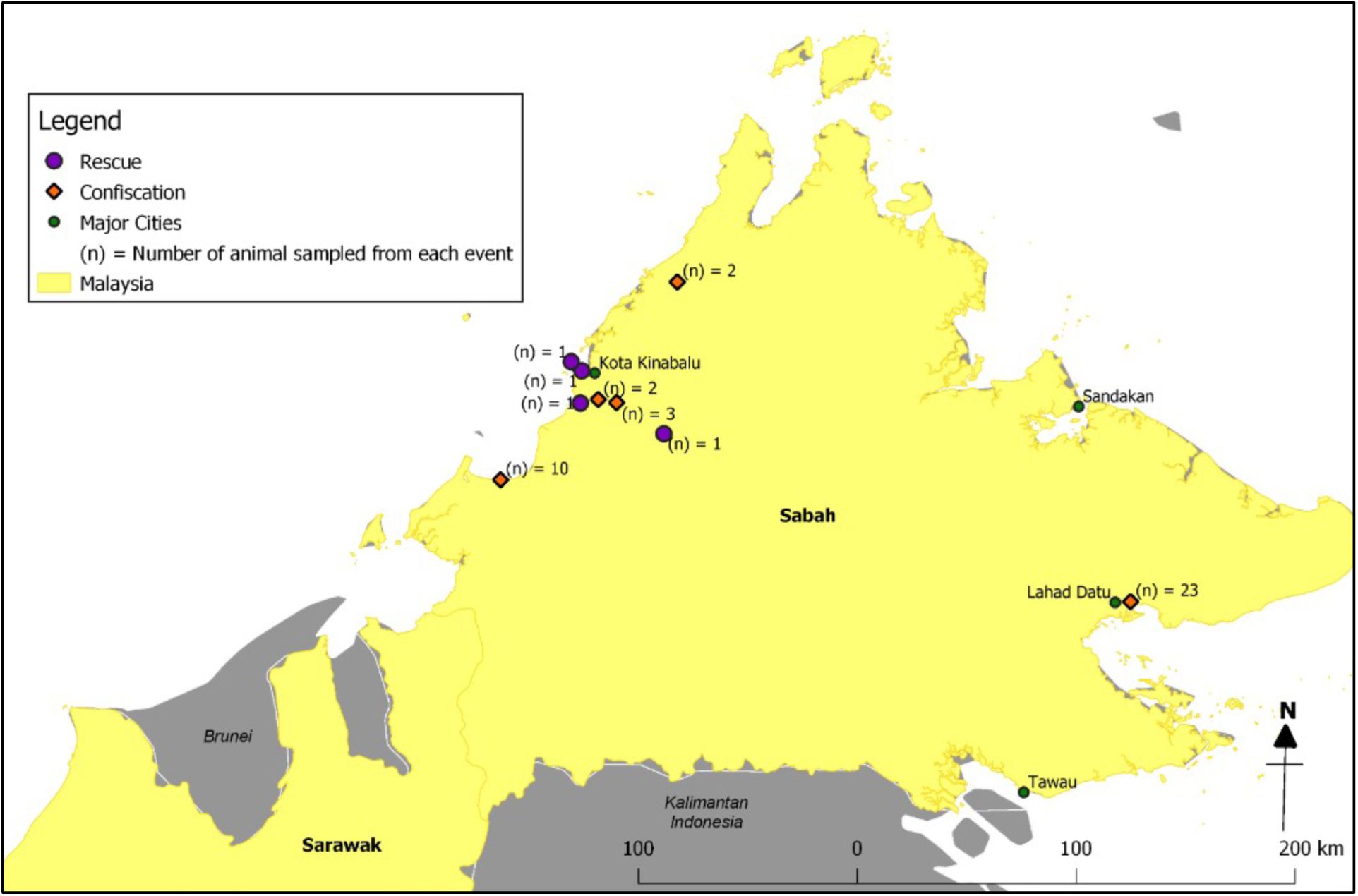
Locations where pangolins were rescued and confiscated in Sabah state.

